# Enhancing Hi-C contact matrices for loop detection with Capricorn, a multi-view diffusion model

**DOI:** 10.1101/2023.10.25.564065

**Authors:** Tangqi Fang, Yifeng Liu, Addie Woicik, Minsi Lu, Anupama Jha, Xiao Wang, Gang Li, Borislav Hristov, Zixuan Liu, Hanwen Xu, William S. Noble, Sheng Wang

**Author notes:** TF, YL, and AW contributed equally to this work.

## Abstract

High-resolution Hi-C contact matrices reveal the detailed three-dimensional architecture of the genome, but high-coverage experimental Hi-C data are expensive to generate. On the other hand, chromatin structure analyses struggle with extremely sparse contact matrices. To address this problem, computational methods to enhance low-coverage contact matrices have been developed, but existing methods are largely based on resolution enhancement methods for natural images and hence often employ models that do not distinguish between biologically meaningful contacts, such as loops, and other stochastic contacts. We present Capricorn, a machine learning model for Hi-C resolution enhancement that incorporates small-scale chromatin features as additional views of the input Hi-C contact matrix and leverages a diffusion probability model backbone to generate a high-coverage matrix. We show that Capricorn outperforms the state of the art in a cross-cell-line setting, improving on existing methods by 17.8% in mean squared error and 22.9% in F1 score for chromatin loop identification from the generated high-coverage data. We also demonstrate that Capricorn performs well in the cross-chromosome setting and cross-chromosome, cross-cell-line setting, improving the downstream loop F1 score by 15.7% relative to existing methods. We further show that our multi-view idea can also be used to improve several existing methods, Hi-CARN and HiCNN, indicating the wide applicability of this approach. Finally, we use DNA sequence to validate discovered loops and find that the fraction of CTCF-supported loops from Capricorn is similar to those identified from the high-coverage data. Capricorn is a powerful Hi-C resolution enhancement method that enables scientists to find chromatin features that cannot be identified in the low-coverage contact matrix. Implementation of Capricorn and source code for reproducing all figures in this paper are available at https://github.com/CHNFTQ/Capricorn.

## 1 Introduction

Chromosomes encode genetic and epigenetic cellular programs, leveraging a complex three-dimensional (3D) genome architecture in eukaryotic cells that is critical for many biological processes, including modulating gene regulatory relationships, RNA splicing sites, and DNA repair mechanisms [3, 8]. These architectures within a cell can be measured with assays such as high-throughput chromosome conformation capture (Hi-C) [27], genome architecture mapping (GAM) [1], split-pool recognition of interactions by tag extension (SPRITE) [34], and HiChIP [31]. *High-resolution* Hi-C datasets employ smaller bin sizes, thereby more precisely characterizing genomic substructures, but consequently require more experimental reads in order to produce a suitably dense matrix. Importantly, doubling the resolution requires quadrupling the experiment read counts due to the pairwise nature of interactions [44].

In order to obtain denser contact matrices, computational *resolution enhancement* methods take low-coverage, high-resolution Hi-C matrices with fewer measured contacts and generate the corresponding highcoverage, high-resolution matrices. Existing approaches adopt techniques from computer vision, such as convolutional neural networks [26, 29, 30, 48] and generative adversarial networks (GANs) [9, 18, 19, 21, 28] to produce detailed contact maps. The resulting data can then be used to perform analyses of genome folding, including classification of large-scale chromatin features like A/B compartments [6, 12, 15, 17, 27] and identification of small-scale chromatin features like topologically associating domains (TADs) [7, 10, 11, 36, 45] and chromatin loops [37, 38, 40, 43, 47]. Although existing approaches aim to minimize mean-squared error (MSE) and perform well according to metrics developed for natural image analyses, these metrics may not be well suited to teach the model to capture biologically relevant chromatin features, and especially small-scale loops.

We hypothesize that resolution enhancement can produce contact matrices that better capture these higher-order chromatin structures if we design a loss function that explicitly models structures like loops and TADs during resolution enhancement (Table 1). Such an approach can both provide additional supervision for the resolution enhancement task and help teach the model to distinguish and enhance particularly interesting genomic contacts. We therefore propose Capricorn, which incorporates additional biological views of the contact matrix to emphasize important chromatin interactions, while leveraging powerful computer vision diffusion models [41] for the model backbone (Figure 1). In particular, Capricorn learns a diffusion model that enhances a five-channel image, containing both the primary Hi-C matrix as well as representations of TADs, loops, and distance-normalized counts computed from the original low-coverage matrix. Capricorn thereby learns meaningful structural contacts as well as the general high-coverage matrix structure.

**Table 1:** A summary of existing Hi-C resolution enhancement methods. We indicate whether each method adopts common computer vision loss terms, with “TV” for total variation and where pixel-wise losses include mean squared error (MSE) and L1 loss. Notably, only VEHiCLE [19] and Capricorn incorporate biological features into the resolution enhancement model. Although VEHiCLE uses TAD identification in its loss, Capricorn is trained to enhance TAD and loop features along with the contact matrix. Capricorn is the only method that uses a diffusion model as the backbone. We limit the comparison to methods for Hi-C resolution enhancement for proximal contacts (interaction within 2 megabases) that do not require additional high-coverage input data, excluding distal-contact enhancement techniques like BoostHiC [5] and reference-based methods like RefHiC-SR [49].

| Method | Publication date | Pixel-wise loss | Adversarial loss | TV loss | Perceptual loss | Biology | Backbone |
| --- | --- | --- | --- | --- | --- | --- | --- |
| HiCPlus [48] | Feb. 2018 | ✓ |  |  |  |  | CNN |
| hicGAN [28] | July 2019 |  | ✓ |  |  |  | CNN (GAN) |
| HiCNN2 [30] | Oct. 2019 | ✓ |  |  |  |  | CNN |
| HiCNN [29] | Nov. 2019 | ✓ |  |  |  |  | CNN |
| DeepHiC [21] | Feb. 2020 | ✓ | ✓ | ✓ | ✓ |  | CNN (GAN) |
| SRHiC [26] | Apr. 2020 | ✓ |  |  |  |  | CNN |
| HiCSR [9] | July 2020 | ✓ | ✓ |  | ✓ |  | CNN (GAN) |
| VEHiCLE [19] | Apr. 2021 | ✓ | ✓ |  |  | ✓ | CNN (GAN) |
| EnHiC [24] | July 2021 | ✓ | ✓ |  | ✓ |  | CNN (GAN) |
| HiCARN-1 [18] | Apr. 2022 | ✓ |  | ✓ | ✓ |  | CNN |
| HiCARN-2 [18] | Apr. 2022 | ✓ | ✓ | ✓ | ✓ |  | CNN (GAN) |
| **Capricorn** | (ours) | ✓ |  |  |  | ✓ | Diffusion |

**Figure 1.**
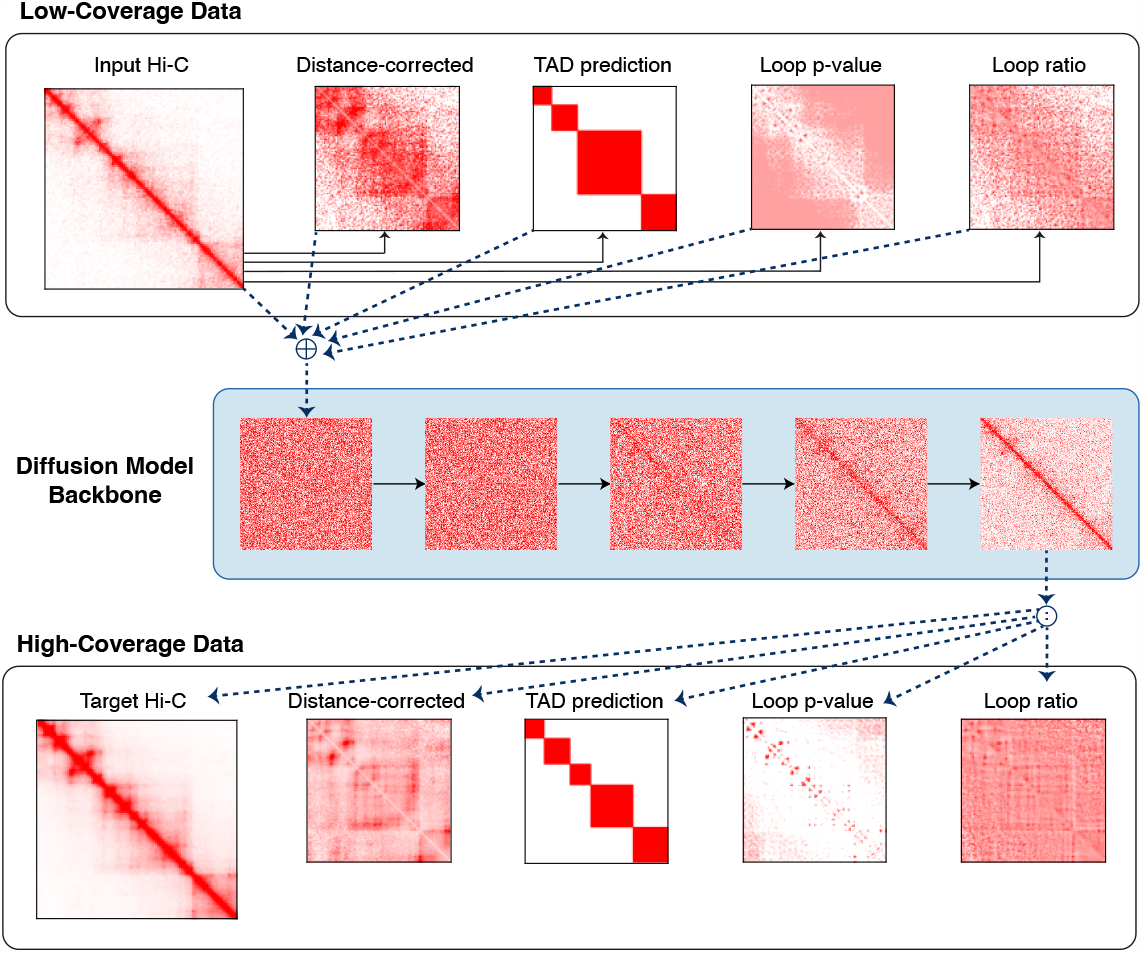
Overview of Capricorn architecture. Given the low- and high-coverage Hi-C contact matrices we compute small-scale chromatin features that explicitly teach Capricorn to recognize biologically meaningful contacts. We then leverage a diffusion model backbone to iteratively de-noise a random contact matrix, conditioned on the low-coverage data.

We compare Capricorn to four existing Hi-C resolution enhancement approaches representing the state of the art, and we find that Capricorn outperforms the others in terms of both MSE and its ability to detect loops from the predicted high-coverage contact matrix. We tested the models’ generalizability across both cell line and chromosome and found that Capricorn’s enhanced matrices had a 17.8% lower MSE and 22.9% higher loop F1-score on average when transferring learned patterns across cell lines. We further found that Capricorn’s key idea of incorporating higher-order chromatin features as additional input views is a broadly applicable technique that improved the comparison approaches as well, though Capricorn’s diffusion model backbone still provides the best performance. As a final validation, we use DNA sequence to evaluate the fraction of identified loops from the high coverage, low coverage, and Capricorn-generated contact matrices that are supported by flanking CTCF motifs, and find that Capricorn’s loops have comparable CTCF support to the high-coverage-derived loops. In summary, Capricorn is a general-purpose approach for chromatin conformation capture contact matrix resolution enhancement, and in the future additional feature views can be easily incorporated into the framework for various downstream tasks of interest.

## 2 Method

### 2.1 Problem setting

For a set of experimental cell lines 𝒞, we have interaction frequency matrices for chromosomes Ξ. In the supervised resolution enhancement task, we are given a dataset with pairs of low- and high-coverage contact matrices 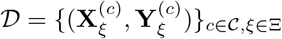, all at high resolution. For chromosome *ξ* containing *N*_*ξ*_ total base pairs and a fixed resolution Δ, the contact matrix shapes are identical, with 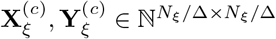. However, the high-coverage version 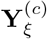 has *λ*-fold more contacts in the matrix.

The aim is to obtain a model *f* such that 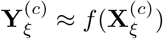 for all *c*∈ 𝒞 and *ξ*∈ Ξ. Furthermore, *f* should generalize to both new cell lines and different chromosomes. Because most Hi-C interactions occur between nearby intrachromosomal loci, we restrict the model to enhance intrachromosomal contacts within 2 Mb of each other as in previous work [18, 28].

Given the matrix 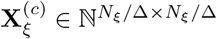 each entry 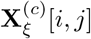 reports a Δ^2^ = 10kb× 10kb region of genomic interactions between genomic regions [10*i* kb, 10(*i* + 1) kb) and [10*j* kb, 10(*j* + 1) kb), where the bin size is defined as Δ = 10 kb. We then tile the paired contact matrices into 40 × 40 non-overlapping submatrices 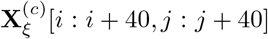, each covering a 400^2^ kb region of contacts, consistent with many Hi-C resolution frameworks [9, 18, 21, 26, 29, 30, 28, 48]. For simplicity of notation, we drop the cell line and chromosome annotation when comparing a pair of low- and high-coverage tiled matrices, and we refer to a pair as (**X, Y**).

### 2.2 Chromatin structure resolution enhancement

The key idea behind Capricorn is that explicitly modeling small-scale chromatin features such as TADs and loops will improve the method’s ability to focus on meaningful contacts. We therefore train Capricorn to enhance the biological interpretation of the low-coverage contact matrix as well as the low-coverage contact matrix itself, thereby explicitly training the model to recognize important 3D chromatin structures from low-coverage data.

Toward this end, in addition to the resolution enhancement task with paired (**X, Y**), we further include additional views of chromatin features derived from **X** and **Y** for resolution enhancement.

1. Distance-corrected: (**X**^(*oe*)^, **Y**^(*oe*)^). The original contact matrices contain a strong bias based on inter-locus distance, with a higher number of measured contacts for locus pairs close to the diagonal. We therefore include a contact matrix corrected for expected contacts based on distance. Specifically, we compute the expected matrix by taking the average along each diagonal

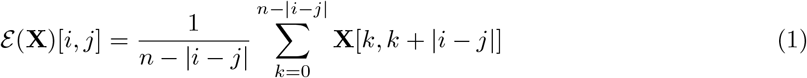

for all *i* = 1, …, *n* and *j* = 1 …, *n*, where *n* = *N*_*ξ*_*/*Δ for notational simplicity. We then normalize the observed matrix, clamp any large values, and finally normalize to the range [0, 1] as

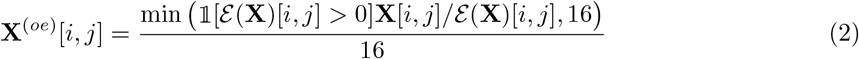

where 𝟙 [·] is the indicator function, and compute **Y**^(*oe*)^ analogously.
2. TAD score: (**X**^(*tad*)^, **Y**^(*tad*)^). We compute the insulation score (IS) [7] for TAD detection, using sliding windows along the contact matrix diagonal to identify insulated regions with low scores and in-domain regions with high scores. We then smooth the IS scores by averaging them over a 21-bin domain and assign the within-TAD label 1 to contiguous regions with monotonically decreasing ISs centered around some locus (*i, j*), where all ISs are still larger than the contact matrix average IS and the IS of (*i, j*) is at least 0.1 larger than the IS at the region boundary.
3. Loop p-value and ratio: (**X**^(*loop*−*p*)^, **Y**^(*loop*−*p*)^), (**X**^(*loop*−*r*)^, **Y**^(*loop*−*r*)^). We follow the Hi-C Computational Unbiased Peak Search (HiCCUPS) loop detection algorithm [37], which tests whether the measured contacts are significantly more frequent than would be expected. The method combines a 10×10 donut kernel centered at a specific locus with a horizontal, vertical, and lower-left quadrant kernel. For a given locus (*i, j*), we use the distance-based expected matrix as defined in (1) to compute

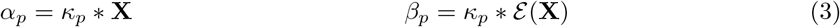

for each kernel *κ*_*p*_ = {*κ*_1_, …, *κ*_4_}. Then we compute the *loop ratio* for the input matrix as

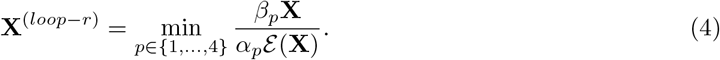

We also follow the HiCCUPS algorithm and fit a Poisson distribution to the observed measurements with *λ*-chunking, where we set *λ* = 2^1*/*3^ as in the default HiCCUPS algorithm [37]. In the *λ*-chunking protocol, we set *μ*_*i,j*_ = max_*c*_ *λ*^*c*^ subject to 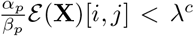, grouping pixels into relative intensity ranges of {[0, 1), [1, 2^1*/*3^), [2^1*/*3^, 2^2*/*3^),…} Finally, we use the result of *λ*-chunking to compute the *loop p-value* as

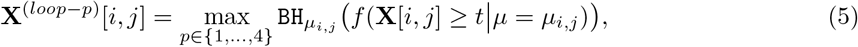

where *f* (± ≥ *t*|*μ*) represents the survival function of the Poisson distribution with expected value *μ*, and 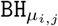 indicates the Benjamini-Hochberg false discovery rate correction [2, 4] over the p-values for all pixels with the same *λ*-chunked *μ*_*i,j*_. We compute **Y**^(*loop*−*r*)^ and **Y**^(*loop*−*p*)^ based on the ground-truth high-coverage matrix **Y** in an analogous manner.

Importantly, we compute each of the low-coverage chromatin features directly from the low-coverage experimental matrix **X**, so the inputs to Capricorn have no knowledge of the high-coverage contact matrix **Y**, which would prevent Capricorn’s practical utility during inference. The original contact matrices and derived chromatin structures are then concatenated to form the full input and output

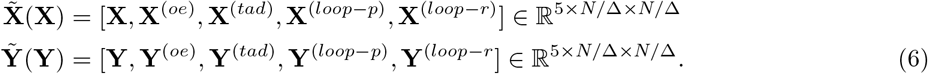

In order to prevent one experimental view from dominating model training, we first trained an exploratory version of Capricorn end-to-end and computed the validation set loss 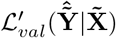. We compared the contribution of each generated view in 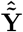 to the overall loss and then computed a weight vector *ω* ∈ ℝ+^5^ that would lead to each view contributing equally to the loss. When training the full version of Capricorn, we therefore weighted each channel by 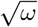 when computing the input and output.

### 2.3 Review of diffusion probability models

To carry out the resolution enhancement task given the low-coverage contact matrix and derived small-scale chromatin features, we use a *diffusion probability model* backbone [20, 46], which has recently been shown to excel in image generation tasks [13, 25, 42] and is much more stable to train than GANs. To the best of our knowledge, Capricorn is the first approach for Hi-C resolution enhancement that leverages diffusion probability models, and we therefore provide a brief overview here.

Diffusion models leverage a latent Markov chain framework that iteratively denoises an input to produce diverse and realistic outputs [20, 46]. The models are split into a *forward process q*(**y**_*t*_|**y**_*t*−1_) and *reverse process p*(**y**_*t*−1_|**y**_*t*_, **x**), where **y**_0_ indicates the original, high-fidelity image, **y**_*t*_ are latent variables with the same dimension as **y**_0_, and **x** is an optional input term on which to condition the process. Specifically, the forward process computes progressively noisier latent representations of the true image as

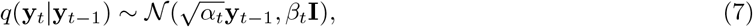

where *β*_*t*_ is defined by a noise schedule, such as a cosine schedule [32, 41], *α*_*t*_ = 1−*β*_*t*_, and the Markov chain repeats for *t* = 1, …, *T*. The reverse diffusion process, optionally conditioned on input **c**, can be then parameterized by a learnable *p*_*θ*_(·) as

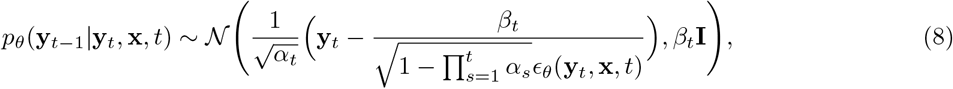

where *ϵ*(·) is trained to predict noise from the previous step’s **y**_*t*_ prediction and the conditional input **x**. If the total noise 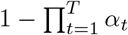 added by all steps in the Markov chain is sufficiently large, then the final noisy representation is well approximated by **y**_*T*_ ∼ 𝒩(**0, I**), enabling inference from a random Gaussian sample.

### 2.4 Low-coverage guided diffusion

We leverage conditional diffusion probability models for the resolution enhancement task. The combined contact matrix and its derived chromatin feature views can be interpreted as a five-channel image, so we approach the problem as high-coverage image generation where low-coverage inputs used to guide the generation process. Specifically, Capricorn is built around a backbone that approximates 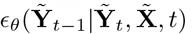 as part of the reverse process given in (8) (Figure 1).

Capricorn’s multi-channel framework encodes additional biological understanding of the contact matrix data into the diffusion model resolution enhancement task, teaching the model to identify significant contacts while enabling the standard MSE loss formulation. Specifically, we optimize the objective

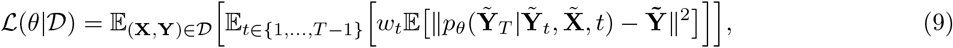

where *w*_*t*_ is a diffusion loss hyperparameter impacting the relative amount of sampled Gaussian noise [41], and the reverse diffusion process begins after sequentially introducing *t* noise samples to the true matrix views 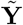 for training.

### 2.5 Performance measures

We evaluate the results using both image-based and biologically motivated metrics, focusing on the generated high-coverage contact matrix. Specifically, given a model’s predicted contact matrix 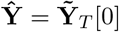, we compute the MSE as 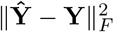, for ∥·∥_*F*_ the Frobenius norm.

We additionally use the enhanced high-coverage matrix to call loops with the HiCCUPS algorithm [37] to compute loop ratios and p-values as described in (4) and (5) respectively. For all loci (*i, j*) with p-value *<* 0.1 and observed-over-expected ratio larger than 1.75 for the donut or lower-left-quadrant kernels, or larger than 1.5 for the horizontal or vertical kernels, we annotate (*i, j*) as containing a loop. We then compute the loop *F* 1 score 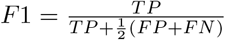 with a 5 pixel tolerance range, such that *TP* are the true positive loops called from the predicted data that appear within [*i*−5 : *i* + 5, *j*−5 : *j* + 5] in the loops called from the ground-truth data, *FP* are the loops called from the predicted data that do not appear within the five-pixel range from the ground-truth data, and *FN* the loops that are called from the ground-truth data but do not occur in the five-pixel tolerance range for predicted data.

### 2.6 Hi-C data preprocessing

We collected Hi-C data using the Gene Expression Omnibus (GEO) database for the GM12878 Epstein-Barrvirus-infected human lymphoblastoid cell line and K562 human chronic myelogenous leukemia lymphoblast cell lines from the Rao et al. (2014) [37] dataset (accession code GSE63525), restricting the contact matrix to read mapping quality≥30 and processed to 10 kilobase (kb) resolution, following previous work [18]. We adopt the contact matrix preprocessing techniques from HiCARN [18] and DeepHiC [21], including tiling the contact matrices into 40×40 submatrices, only retaining submatrices in the 2 megabase (Mb) region around the diagonal, clamping the high-coverage matrix to [0, 255] and then normalizing to [0, 1], and clamping the low-coverage matrix to [0, 100] and then normalizing to [0, 1].

In order to simulate low-coverage data, we randomly downsampled the GM12878 and K562 cell line Hi-C matrices [37] to 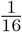 of the original read count. We treated these downsampled data as the low-coverage matrices and the original matrices as their full-coverage pairs. We conduct and report on two separate experiments to evaluate model generalizability: in the first, we train on the GM12878 data and test the model on K562; in the second, we train on K562 data and test on GM12878. In both experiments, we withhold chromosomes 4, 5, 11, and 14 from the training cell line as our validation set. The K562 data is excluded for chromosome 9 due to extreme sparsity at 10 kb resolution. After running the exploratory version of Capricorn to determine view weighting, as described in Section 2.2, we set view weights to 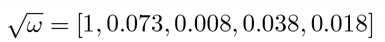 when training on GM12878 and 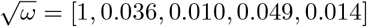 when training on K562.

### 2.7 Existing model implementations

We use the publicly available python HiCARN [18] repository from GitHub (https://github.com/OluwadareLab/HiCARN) for both their model and contact matrix preprocessing implementation, as well as source code implementations for HiCNN [29] and HiCSR [9].

We also use the conditional diffusion probability model Imagen [41] as the resolution enhancement backbone model, updating the model to condition on low-coverage contact matrices rather than text. We choose Imagen rather than other image diffusion models [33, 35, 39] due to its efficient U-Net architecture, which is faster and more memory efficient than other diffusion generators. We use the GitHub repository https://github.com/lucidrains/imagen-pytorch to access Imagen. This enabled model training in approximately 28 hours and inference in approximately 45 minutes on an NVIDIA A4000 GPU in the cross-cell-line experiment.

## 3 Results

### 3.2 Capricorn accurately enhances contact matrices and loop features

We first sought to evaluate Capricorn in the cross-cell-line setting, where the model is trained on the simulated low-coverage data and measured high-coverage data on one cell line and tested on the simulated low-coverage data of another cell line. We find that Capricorn outperforms other methods both in terms of its ability to enhance chromatin loops from the low-coverage data and its accuracy in producing high-coverage data for both test cell lines in the cross-cell-line experimental setting. We first used the HiCCUPS [37] algorithm to call loops in the resolution-enhanced contact matrices, and we compared these to the loops HiCCUPS identified in the original full-coverage data. As shown in Figure 2a,b, Capricorn outperformed all other approaches in its ability to recognize and enhance loop chromatin structures when transferred to both the GM12878 and K562 cell lines, with an average loop *F*_1_ score taken over chromosomes of 0.48 on GM12878 test data and 0.31 on K562 test data, relative to 0.43 and 0.20 for HiCSR, the best-performing comparison approach, and 0.30 and 0.19 if directly using the input low-coverage data scaled by the experimental downsampling rate. The performance difference for all methods between GM12878 and K562 test data can be explained by the difference in pairwise contacts in the original dataset: GM12878 has approximately five times more measured contacts than K562 [37]. These results demonstrate that Capricorn is able to successfully identify and enhance meaningful biological contacts, such as those involved in loop formation.

**Figure 2.**
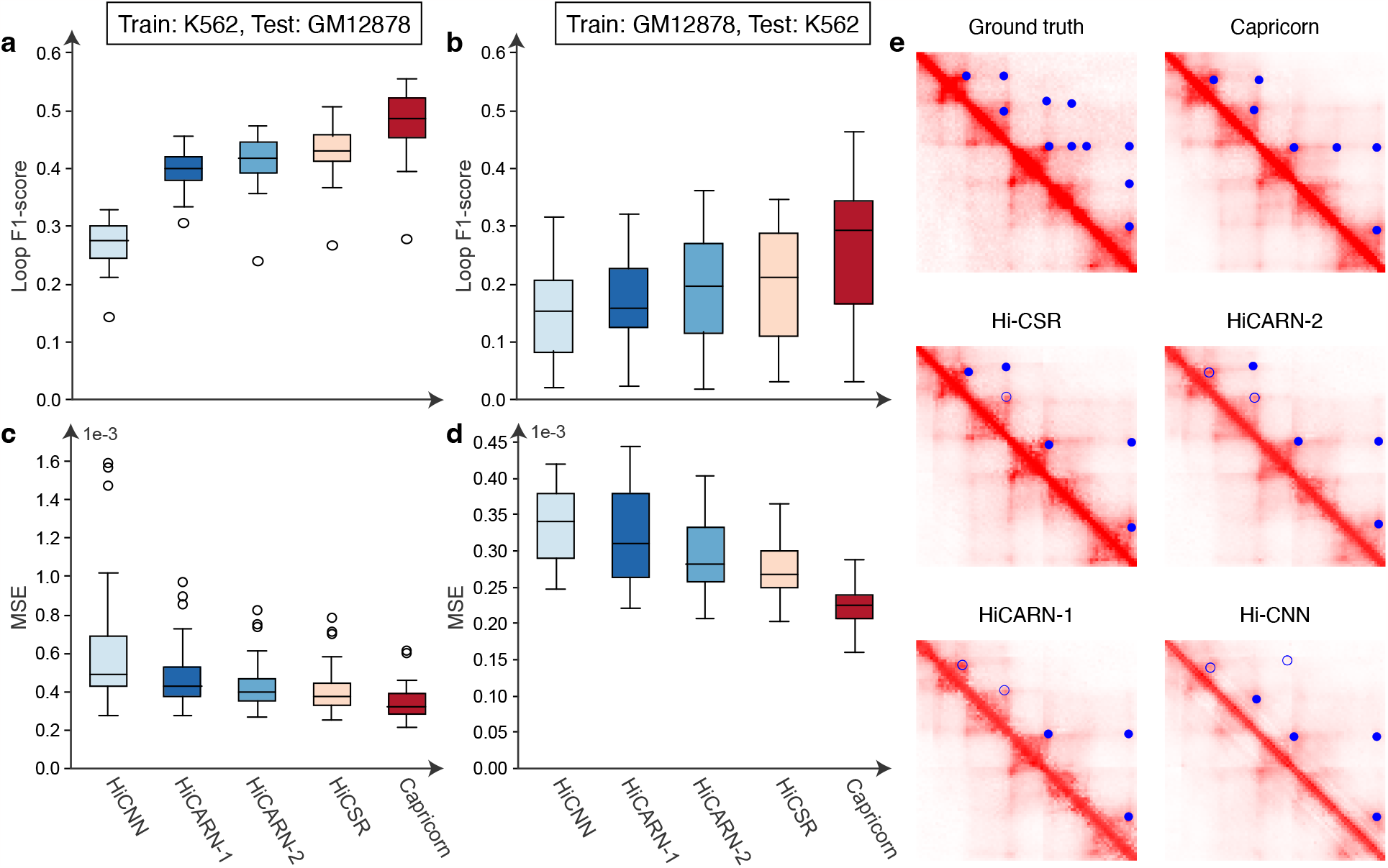
Resolution enhancement model performance. **a-d**, Boxplot comparison of F1-score for loop detection from the generated high-coverage matrices (**a**,**b**, higher is better) and generated matrix MSE (**c**,**d**, lower is better). The boxplots show the median, interquartile range (IQR), 1.5 ×IQR, and outliers by chromosome. **e**, Ground-truth high-coverage submatrix covering genomic loci from 47.3 Mb to 48.1 Mb on GM12878 chromosome 17, compared to the generated high-coverage submatrix output by each method. Blue circles indicated called loops in the ground-truth or predicted high-coverage data. Circles for loops that are also called in the ground truth data are filled in; loops that are called from the generated data but not the ground-truth data are empty circles.

Furthermore, Capricorn achieved a lower prediction MSE than any of the comparison approaches, indicating that the additional views can also boost the overall resolution enhancement results. Figure 2c,d, shows that Capricorn has an average MSE taken over chromosomes of 3.5e−4 and 2.2e−4 for the GM12878 and K562 test cell lines, respectively, relative to 4.2e−4 and 2.8e−4 for HiCSR, again the best-performing comparison approach, and to 2.4e−3 and 1.1e−3 if directly using the input low-coverage data scaled by the experimental downsampling rate.

### 3.2 Small-scale chromatin features are critical to model improvement and are model-agnostic

To better attribute Capricorn’s strong performance, we next investigated the specific impact of including small-scale chromatin features as additional views to train the resolution enhancement model and enhance structurally meaningful contacts. We compared Capricorn’s performance when trained to enhance chromatin features as well as the Hi-C matrix to its performance when performing resolution enhancement without any additional views. As shown in Figure 3a,b, we find that explicitly training the model to enhance meaningful biological contacts, as captured in the small-scale chromatin features, significantly improves our ability to identify these features from the enhanced contact matrices (one-sided Wilcoxon signed-rank test p-value *<* 6.4e−5, *<* 1.3e−6 comparing loop F1-score over submatrices for test GM12878 and K562 respectively).

**Figure 3.**
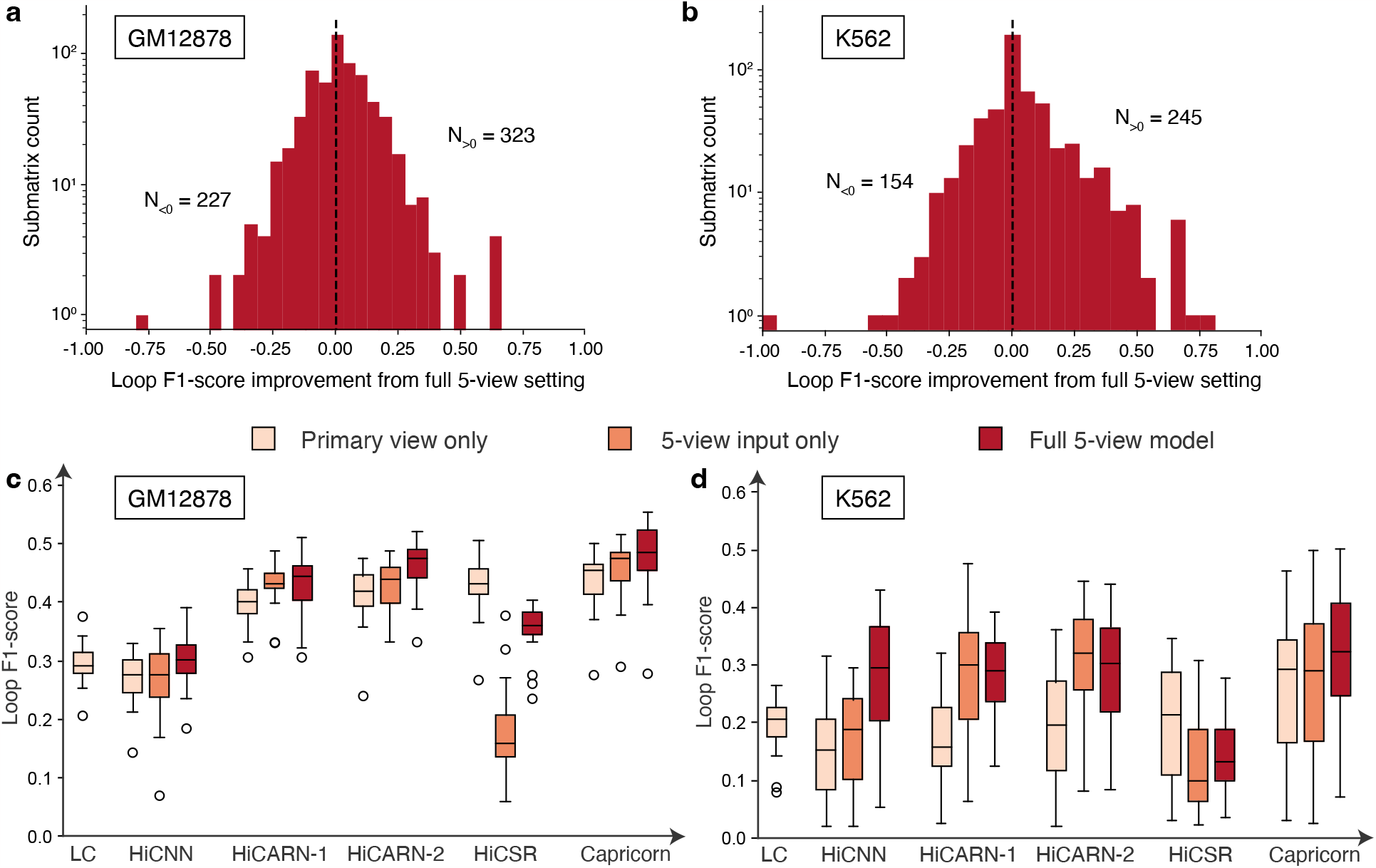
Study of Capricorn’s multi-view chromatin feature framework. **a**,**b**, Histogram of the difference in loop F1-score from the predicted high-coverage contact matrix in the GM12878 (**a**) and K562 (**b**) test datasets using the full five-view Capricorn framework and an alternate version of Capricorn that only includes the primary Hi-C matrix view as the input and output for the diffusion model backbone. The counts *N*_*>*0_ and *N*_*<*0_ indicate the number of submatrices for which the five-view setting respectively outperform or underperform the primary-only setting. **c**,**d** Boxplot comparing the performance of resolution enhancement methods in the primary-view-only setting where enhancement is only performed with Hi-C matrices, the five-view-input setting where small-scale chromatin features are included as input to the resolution model, and the full-five-view setting where small-scale chromatin features are used both as input and output to train the model backbone. “LC” indicates the results taking low-coverage matrix and scaling all contacts by the downsampling factor 16. The boxplots show the median, IQR, 1.5×IQR, and outliers by predicted submatrix.

Because our multi-view idea is not architecture dependent, we then tested whether the benefits of including small-scale chromatin features as part of model training generalized to other resolution enhancement network architectures. To this end, we updated each of the comparison models to accept multi-channel input matrices and compared the results of three model formulations (Figure 3c,d).

1. Primary view: This setting uses the comparison models’ default resolution enhancement pipelines with the low- and high-coverage Hi-C matrices as input and output. We also consider Capricorn’s performance when trained without the additional biological views in this setting.
2. Five-view input only: This setting uses the additional chromatin feature views as input to the model, but is still trained to only predict the high-coverage view. As shown in Figure 3c,d, many methods perform better in this setting than the original primary view, indicating the utility of the additional biological feature inputs.
3. Full five-view model: This setting uses Capricorn’s complete multi-view setting, including all five chromatin feature views as input and training the model to enhance the small-scale chromatin features in addition to the Hi-C matrices. This setting yields the best downstream loop calling performance for Capricorn and three of the four comparison approaches. Although the multi-view biological inputs from the five-view input only setting generally improve performance relative to the primary view only setting, the additional performance gains in this setting highlight the additional value of including small-scale chromatin features as model outputs that are explicitly included in the loss function.

Although we show our key multi-view idea to be generalizable to many model architectures and loss formulations, we still find that Capricorn’s diffusion model backbone outperforms the convolutional models. Comparing the full five-view model results for Capricorn and HiCARN-2, the best-performing comparison approach, Capricorn still outperforms the other approaches on both GM12878 and K562 test data (one-sided Wilcoxon signed-rank p-value *<* 7.5e−3 and *<* 2.1e−3 respectively).

Capricorn’s multi-view framework is beneficial for loop calling in the generated high-coverage matrix for all models except HiCSR. This observation points to the broad applicability of Capricorn’s key idea, and also suggests opportunities for including additional Hi-C-derived views based on expert domain knowledge for various downstream genome folding analyses. We hypothesize that HiCSR performs poorly when including additional biological views due to its denoising autoencoder component [9], which distinguishes its architecture from HiCARN-2’s convolutional GAN architecture. HiCSR trains this denoising autoencoder based on the setting outputs in order to model a perceptual loss term that is specific to Hi-C matrices. However, this approach may require further hyperparameter tuning such that the perceptual loss term is well balanced with components of the overall loss function, particularly in the full five-view model setting where the additional biological views are modeled by the denoising autoencoder.

### 3.3 Capricorn generalizes across chromosomes

To confirm that Capricorn’s strong performance is not due to memorizing the training chromosomes with relatively small cell-line differences, we conducted a second experiment to rigorously test Capricorn in the cross-chromosome setting. Here, we withhold chromosomes 2, 6, 10, and 12 as a validation set and reserve chromosomes 4, 14, 16, and 20 as test data. We therefore carried out two experiments to examine cross-chromosome and intra-cell-line generalization as well as cross-chromosome and cross-cell-line generalization.

We find that Capricorn is able to transfer its learned resolution enhancement patterns to never-before-seen genomic loci in the cross-chromosome intra-cell-line setting better than other methods (one-sided Wilcoxon signed-rank test p-value *<* 4.0e−3). Across the four test chromosomes, Capricorn’s generated high-coverage contact matrix had an average loop F1-score of 0.55 in the GM12878 experiment (Figure 4a) and 0.34 in the K562 experiment (Figure 4b), relative to the best-performing comparison approaches at 0.49 for HiCARN-1 and 0.25 for HiCARN-2. Similarly, Capricorn had the highest loop F1 scores in the cross-chromosome, cross-cell-line experiments as shown in Figure 4c,d (one-sided Wilcoxon signed-rank test p-value *<* 1.9e−2), with an average loop F1 score of 0.48 on GM12878 and 0.22 on K562 test data, relative to the best comparison approaches respective loop F1-scores of 0.32 from HiCARN-2 and 0.15 from HiCARN-1. As in the cross-cell-line setting, all methods perform better on GM12878 test data because it contains many more measured contacts than the K562 experimental data. This result highlights Capricorn’s ability to generalize the informative, small-scale chromatin patterns it learns across genomic loci as well as cell lines, reiterating the effectiveness of diffusion-based modeling of additional chromatin feature views.

**Figure 4.**
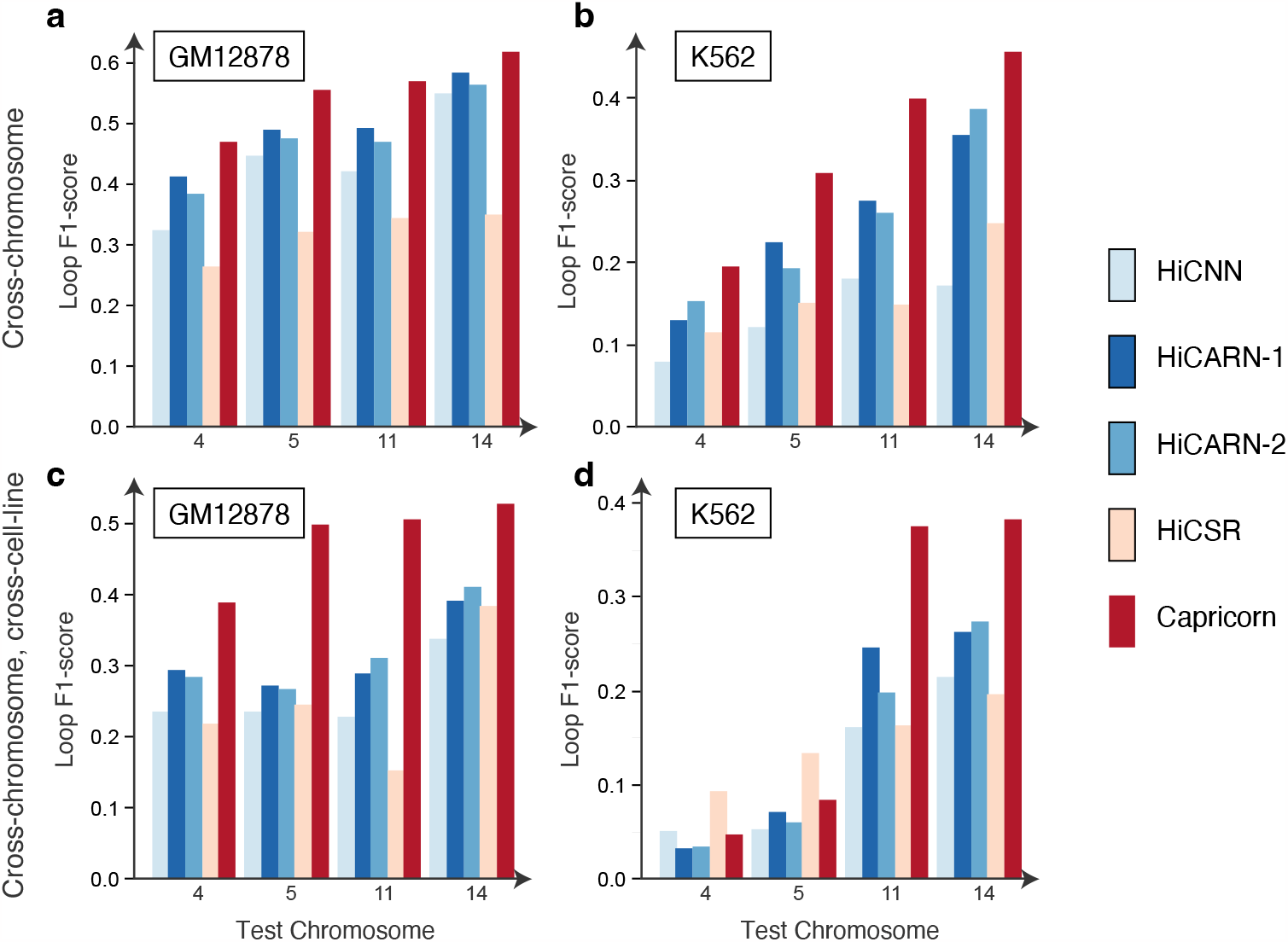
Model performance comparison in cross-chromosome and cross-chromosome, cross-cell-type experimental settings. **a-d**, Barplots showing loop F1-score by test chromosome in a cross-chromosome experiment (**a**,**b**) and cross-chromosome, cross-cell-type experiment (**c**,**d**). Plots are labeled with the cell line of the test data.

### 3.4 Loops discovered with Capricorn are enriched for CTCF motifs

Finally, we leveraged DNA sequence information to further validate the loops identified from Capricorn’s generated high-coverage contact matrices. The CCCCTC-binding factor (CTCF) is a key protein in 3D structure determination for mammalian genomes [16], and many loops anchor at CTCF motifs [37]. In particular, cohesin-mediated loop formation is facilitated by pairs of flanking CTCF motifs occurring in opposite orientations. Because our methods do not make use of the primary DNA sequence, we can use pairs of inward-facing CTCF motifs as additional experimental evidence in support of a candidate loop.

Accordingly, we designed an experiment to compare the extent to which loops discovered based on the high-coverage matrix, the low-coverage matrix, and the Capricorn-enhanced matrix are supported by paired CTCF motifs. We used FIMO [14] to search the human reference genome (build hg38) for loci containing a CTCF motif, using a consensus sequence motif and a p-value threshold of 1e−4, following previous work [37]. This analysis yielded 735,079 CTCF occurrences. For each cell line, we called loops from three matrices: the original high-coverage contact matrix, the downsampled matrix, and Capricorn’s high-coverage generated matrix in the cross-cell-line setting. For each loop, we asked whether it was flanked by two CTCF motifs on opposite strands and in inward-facing orientation within 10 kb on either side. Such loops were marked as “CTCF validated.”

This analysis shows that the Capricorn-enhanced matrix identifies plausible loops. We segregated loops into categories based on whether they were detected in various combinations of the Hi-C matrices, and we calculated the proportion of CTCF validated loops in each category (Figure 5). Loops detected from the Capricorn-enhanced data have comparable CTCF support to loops called from the high-resolution matrix, while non-loop loci genome-wide have an average CTCF rate of 1.2%. In the GM12878 test cell line for instance, 2.1% of the loops identified from Capricorn’s enhanced contact matrix that were not called from either the high coverage or low coverage data had nearby inward-facing CTCF motifs, compared to 2.0% of loops called from the high-resolution data. While fewer of Capricorn’s uniquely identified loops in K562 have supporting CTCF motifs within 10 kb, we highlight that the intersection of Capricorn’s loop set with loops identified in either high- or low coverage data has a very high CTCF support, resulting in CTCF support for 2.6% of called loops from Capricorn overall. This analysis therefore increases our confidence that Capricorn is generating data corresponding to plausible loops, even though DNA sequence is not used during the resolution enhancement task.

**Figure 5.**
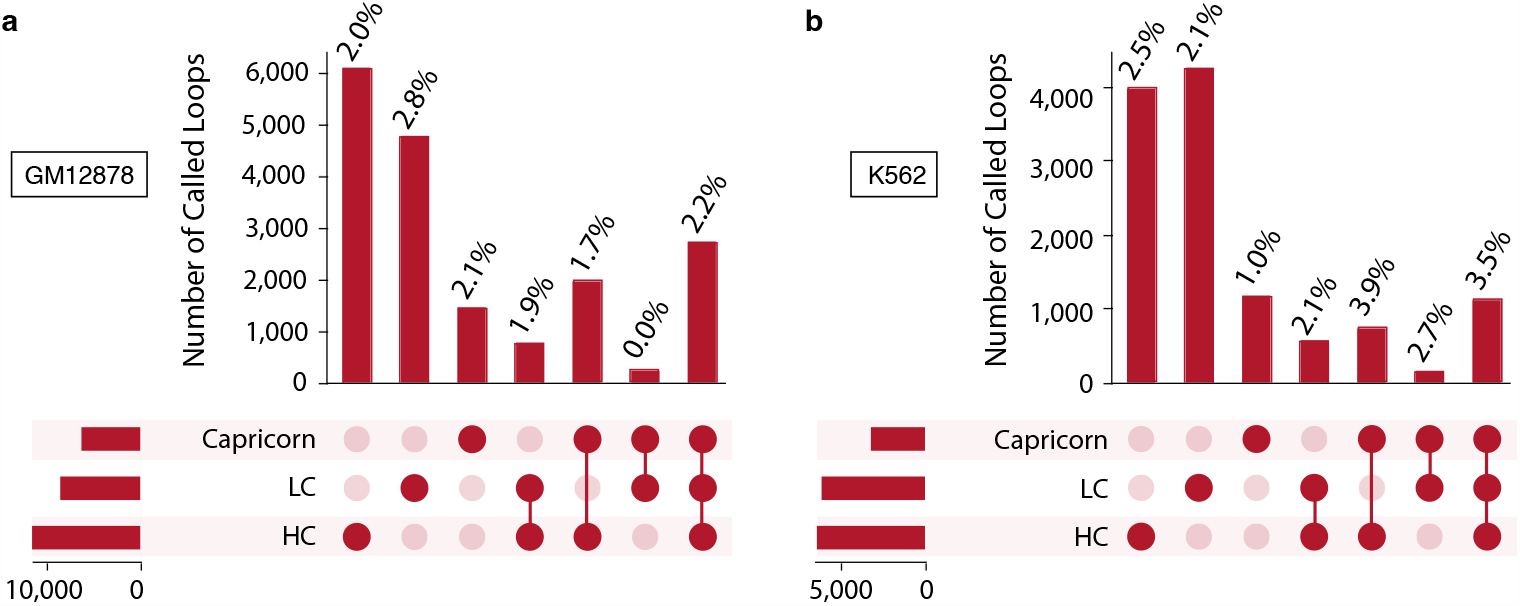
CTCF motif validation of loop calls. **a**,**b** Loops were grouped based on whether they were identified using the high-coverage matrix (“HC”), the low-coverage matrix (“LC”), the Capricorn-enhanced matrix, or some combination thereof in the cross-cell-line experiments for GM12878 (**a**) and K562 (**b**). Two loops were considered the same if both pairs of anchors were no more than 50 kb apart. For each subset of loops, the upset plot indicates the number of loops in that category, with the fraction that are validated by CTCF motifs indicated. For comparison, the fraction of non-loop loci with supporting CTCF motifs was 1.2%.

## 4 Discussion

In this work we present Capricorn as a tool for Hi-C resolution enhancement. Capricorn explicitly models the biology underlying the contact matrices by incorporating small-scale chromatin features into the model formulation and loss function. Furthermore, we find that this key insight is widely applicable to other resolution enhancement approaches, improving performance in three out of four of the comparison approaches. However, the small-scale chromatin feature views still perform best with Capricorn’s conditional diffusion model backbone.

We demonstrate Capricorn’s strong performance in cross-cell-line, cross-chromosome, and cross-chromosome-cross-cell-line settings with the GM12878 and K562 datasets. In all three measured settings, Capricorn is best able to generate high-coverage, high-resolution data containing accurate chromatin loops that can be called with standard tools [37]. This highlights Capricorn’s generalizability as well as the value of including additional biological data views, differentiating Hi-C resolution enhancement from resolution enhancement in natural image applications. Finally, we leverage DNA sequence to further validate the loops identified from Capricorn’s generated data, suggesting that even loops not called from high-coverage data may be reasonable.

In the future, Capricorn’s framework can be further broadened to include additional biological views designed specifically for downstream tasks of interest. In particular, new views that contain structural information covering a more than 400^2^kb locus could help enhancement for TADs and A/B compartments. We also imagine that future work will study the impact of even more model backbones on the multi-view resolution enhancement problem.

Future work can further study Capricorn’s generalizability. Here, we have focused on two human cell lines measured with *in situ* Hi-C. Follow-up work can apply Capricorn to more available Hi-C cell line data and study the impact of training data size. This work could also apply Capricorn to a cross-species transfer learning setting, such as applying the model to mouse data after training only on human data. Another generalizability study could apply a model trained on *in situ* Hi-C data to other contact map types, such as micro-C [23, 22]. Finally, while we leveraged the HiCCUPS [37] loop calling algorithm in this work, future work can test performance across a series of different loop calling methods both for generated high-coverage matrix evaluation and for loop-based biological view construction.

